# From Dataset Curation to Unified Evaluation: Revisiting Structure Prediction Benchmarks with PXMeter

**DOI:** 10.1101/2025.07.17.664878

**Authors:** Wenzhi Ma, Zhenyu Liu, Jincai Yang, Chan Lu, Hanyu Zhang, Wenzhi Xiao

## Abstract

Recent advances in deep learning have significantly improved the accuracy of structure prediction for biomolecular complexes; however, robust evaluation of these models remains a major challenge. We introduce **PXMeter**, an open-source toolkit that support consistent and reproducible evaluation of diverse predictive models across a broad spectrum of biological complex structures. PXMeter provides a unified and reproducible benchmarking framework, offering valuable insights to support the ongoing improvement of structure prediction methods. We also present **a high-quality benchmark dataset** curated from recently deposited structures in the Protein Data Bank (PDB). These entries are manually reviewed to exclude non-biological interactions, ensuring reliable evaluation. Using these resources, we conducted **a comprehensive benchmark** of several structure prediction models, namely Chai-1, Boltz-1, and Protenix. Our benchmarking results demonstrate the advancements achieved by deep learning models, while also identifying ongoing challenges—especially in modeling protein-protein and protein-RNA interactions.

**Project Page:** https://github.com/bytedance/PXMeter

## 1 Introduction

The rapid development of deep learning-based methods has transformed biomolecular structure prediction, enabling high-accuracy modeling of complexes involving proteins, nucleic acids, small molecules, ions and modified residues [1]. However, evaluating these models in realistic and diverse biological scenarios remains a major challenge.

The rigorous assessment of models depends on a reliable and comprehensive evaluation toolkit and high-quality benchmark datasets. Most existing evaluation tools [2–5] either provide only a subset of desired metrics, lack robust support for the outputs produced by modern AI models, or are difficult to extend to incorporate additional models or evaluation metrics. Furthermore, the benchmark dataset collections in common use [6–9] are curated with narrowly defined task-specific objectives, and consequently fail to capture the full diversity of biomolecular interactions. While the AlphaFold 3 paper proposed an evaluation strategy, it did not release the corresponding PDB entries, limiting reproducibility and comparison.

To address these limitations, we introduce PXMeter, an open-source evaluation toolkit that integrates multiple quality checks and supports consistent, reproducible assessment across diverse prediction tasks. We also present a curated benchmark dataset derived from recently deposited structures in the Protein Data Bank (PDB) [10], filtered to exclude experimental artifacts. Using these tools, we conduct a large-scale benchmark of state-of-the-art models—including Chai-1 [11], Boltz-1 [12], and Protenix [13]—across various biomolecular complex prediction tasks. Our contributions are summarized as follows:

- **PXMeter toolkit**. An open-source toolkit for reproducible assessment of complex biomolecular structures—including those with non-standard residues and diverse molecular types—released under the Apache 2.0 License.
- **High-quality benchmark dataset**. A curated dataset for biological complex structure prediction, manually validated to ensure data quality and biological relevance.
- **Model benchmarking**. A comprehensive benchmark and comparative analysis of state-of-the-art prediction models across diverse biomolecular systems, covering both individual molecules and multi-component complexes.

## 2 Results

### 2.1 Development of PXMeter Toolkit

Reliable benchmarking of structure prediction models requires a unified workflow that (i) supports diverse output formats from different models, (ii) establishes atom-to-atom correspondence between predicted and experimental complexes, (iii) reports quality metrics at different granularity, i.e. complex, chain and interface. Most existing tools address only a subset of these needs, often requiring researchers to assemble ad hoc workflows from multiple tools, each with its own process rules-for example, how chains are mapped between ground-truth and prediction, how missing atoms are handled, or how alternative locations are resolved. Such inconsistencies can lead to evaluation bias and reduce the reliability of cross-model comparisons.

To address these challenges, we developed PXMeter, an easy-to-use, end-to-end framework that runs with simple commands. Starting from raw CIF files, PXMeter performs rigorous atom mapping and chain permutation, and generates a unified report that includes DockQ [2], global and interface Local Distance Difference Test (LDDT) [14], and RMSD metrics (Figure 1A). A dedicated mapping engine ensures that every atom evaluated has a corresponding match in the reference structure, preventing inaccurate score inflation due to mismatched identifiers. The package includes a complete data pipeline, from importing raw CIF files and building curated benchmark sets to generating publication-ready results. Its modular codebase allows easy extension, enabling users to incorporate additional metrics or support new models on top of the core mapping engine. Together, these features make PXMeter a reproducible, extensible, and user-friendly tool for fair and consistent evaluation of structure prediction methods.

**Figure 1.**
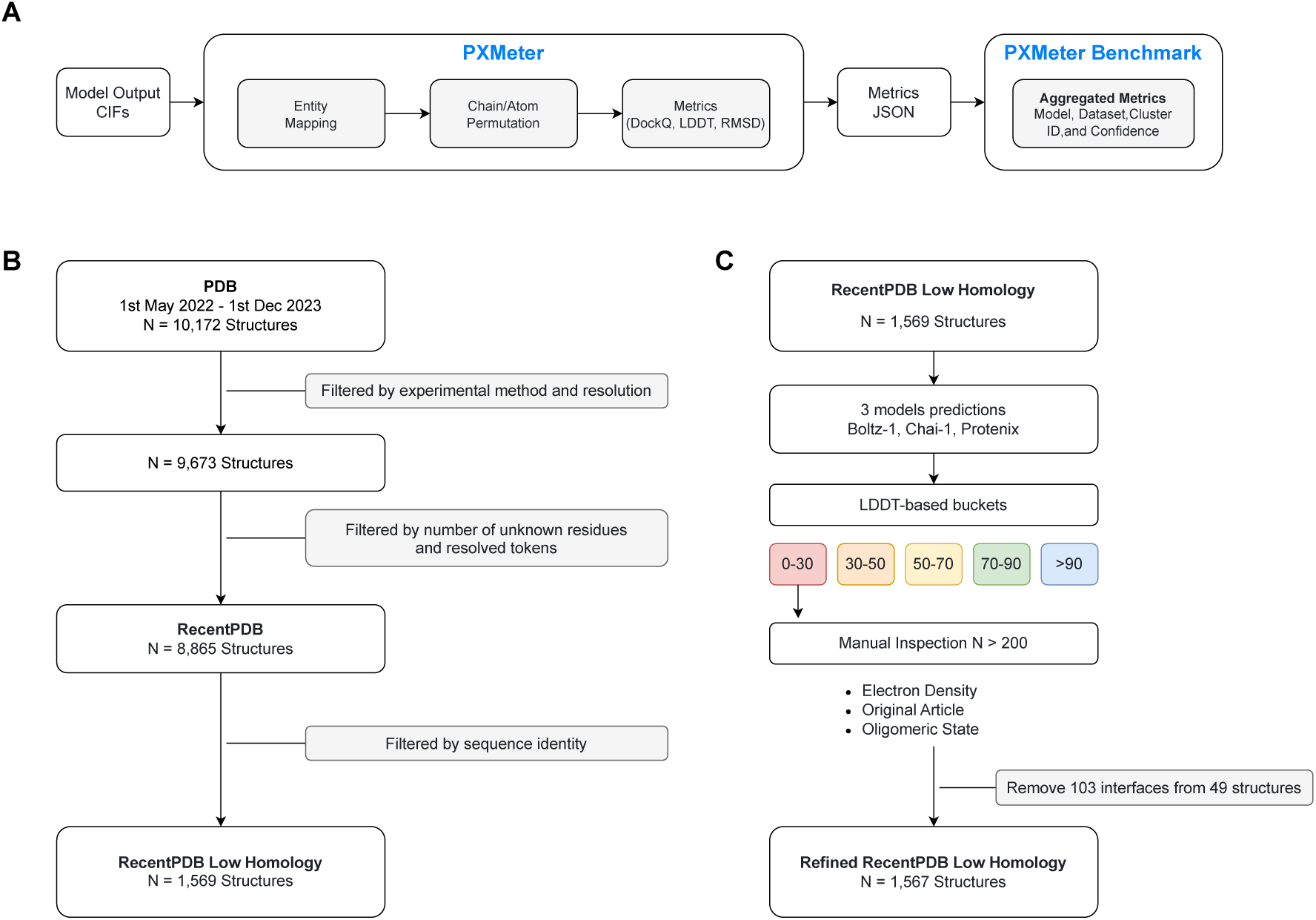
PXMeter workflow and dataset curation process. **A**. Overview of the PXMeter evaluation pipeline. **B**. Construction of the RecentPDB Low Homology sets using combined structural and sequence-based filters. **C**. Manual inspection of samples to exclude structures with experimental artifacts, annotation errors, or related data issues.

### 2.2 Initial Dataset Curation

Starting with the raw mmCIF archives retrieved from the RCSB PDB, we carried out a systematic preprocessing workflow (Figure 1B). Entries were first filtered by deposition date, then screened against a set of compatibility criteria, excluding any structures that could not be processed by baseline models—such as chains with all residues unknown or systems exceeding acceptable size limits. From this curated pool, we constructed the RecentPDB Low Homology dataset, designed to closely replicate the evaluation setup described in the AlphaFold 3 (AF3) study.

To enable more targeted assessments, the curated datasets were further divided by molecular composition and evaluation purpose, resulting in subsets such as protein–protein, protein monomer, antibody–antigen, dsDNA–protein, and RNA–protein (see Section 5.2 for details). For protein–ligand systems, we evaluated model performance separately using the PoseBusters V2 benchmark. All curated structures, together with the metadata and scripts necessary to reproduce all results reported herein, has been made publicly available.

### 2.3 Model Benchmarking Results

#### 2.3.1 Overall Prediction Accuracy Across Complex Types

Based on the curated datasets described above, we next performed a systematic evaluation of three state-of-the-art structure prediction models—Chai-1, Boltz-1, and Protenix—using PXMeter and a set of standardized evaluation metrics. Model performance was assessed using DockQ success rate (> 0.23), interface LDDT (iLDDT) for multimeric interfaces, and pocket-aligned ligand RMSD (< 2 Å) for protein-ligand, as detailed in Section 5.1. For each model, we report three tiers of performance: the Oracle score, representing the best chain/interface structure in the generated ensemble and thereby providing an upper bound on attainable accuracy; the Median score, which approximates the outcome of naive random selection; and the Selected score, corresponding to the top-ranked structure returned by the model’s confidence score (“*ranking_score*” for Protenix; “*aggregate_score*” for Chai-1; “*confidence_score*” for Boltz-1).

##### Protein–ligand docking

As shown in Figure 2A, on the PoseBusters-V2 benchmark, Protenix achieved the highest oracle ligand RMSD success rate (0.910) and maintained a robust 0.833 success rate after selection, outperforming the second best model, Boltz-1, by a significant margin (0.081). This highlights its ability to capture protein-ligand interactions.

**Figure 2.**
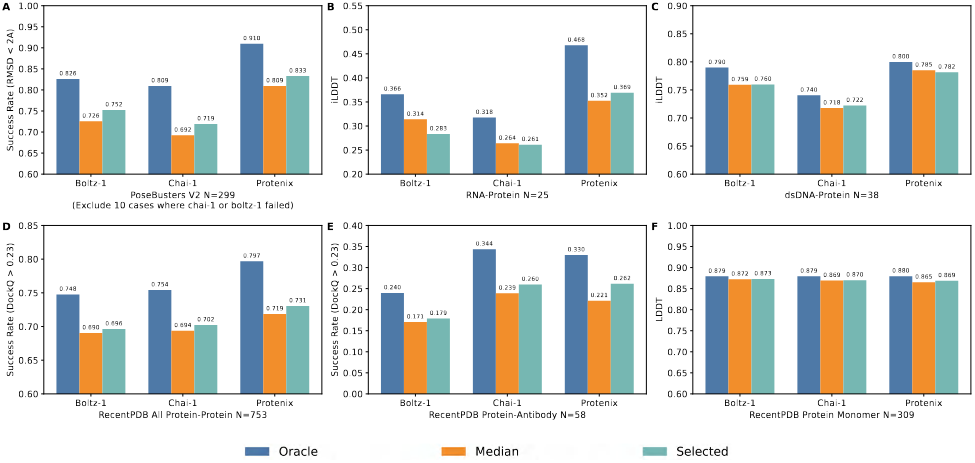
Model performance. Performance comparison of Boltz-1, Chai-1, and Protenix on the curated benchmark datasets detailed in Section 5.2. <u>Oracle</u>, <u>Median</u>, and <u>Selected</u> report metrics for (i) the best chain/interface structure available, (ii) the median chain/interface structure across all generated candidates, and (iii) the top-ranked structure according to the model’s confidence score, respectively. Figure S4 shows the same data with consistent Y-axis scales for comparison. **A**. pocket-aligned ligand RMSD success rates (<u>N</u> denotes the number of targets). Targets lacking predictions from any model were removed, hence the results exclude 10 PDB entries where Boltz-1 or Chai-1 failed. **B, C**. Interface LDDT for protein–nucleic acid complexes (<u>N</u> denotes the number of structures). **D, E**. DockQ success rates for protein–protein and protein–antibody interfaces (<u>N</u> denotes the number of clusters). **F**. Complex LDDT for protein monomers. (<u>N</u> denotes the number of clusters).

##### Protein-nucleic acid interactions

Accurate modeling of protein–nucleic acid complexes remains inherently difficult, due to both the conformational flexibility of nucleic acid backbones [15, 16] and their underrepresentation in training datasets. On the RNA-protein benchmark (Figure 2B), all models showed modest oracle iLDDT values (0.318–0.488). Both Boltz-1 and Chai-1 exhibited limited ranking performance, as indicated by the drop in the selected scores compared to their medians. A similar pattern was observed for Protenix on the AF3 dsDNA-protein subset (Figure 2C), where the selected iLDDT score (0.782) was slightly below the median value (0.785).

##### Protein–protein interactions

Evaluations on the RecentPDB protein–protein benchmarks (Figure 2D, E) show that Protenix achieved the highest DockQ success rates at both the oracle (0.797) and selected (0.731) levels. Protein–antibody interfaces remain particularly challenging due to the high variability of the CDR loops [17, 18], resulting in a marked performance drop across all three models. Within this subset, Chai-1 yielded the highest oracle score, but its selected performance fell slightly behind Protenix, suggesting less effective ranking.

##### Protein monomeric structures

Structure prediction for protein monomers appears comparatively less challenging. On the RecentPDB protein monomer subset, all models reached high LDDT scores (0.865–0.880), with minimal performance drop between oracle and selected tiers.

To verify the reliability of our results, we cross-validated the LDDT and DockQ metrics produced by PXMeter against values reported by OST for a subset of entries (Appendix C), confirming a high degree of consistency between the two tools.

#### 2.3.2 Correlation Between Confidence Score and Structural Accuracy

To assess the effectiveness of the internal confidence scores used by each model, we examined their correlation with actual structural quality across both monomeric and multimeric predictions (Figure 3). For each structure or interface, we selected the top-ranked prediction from 5 seeds ×5 samples and computed Pearson correlation coefficients (PCC) between model confidence and evaluation metrics.

**Figure 3.**
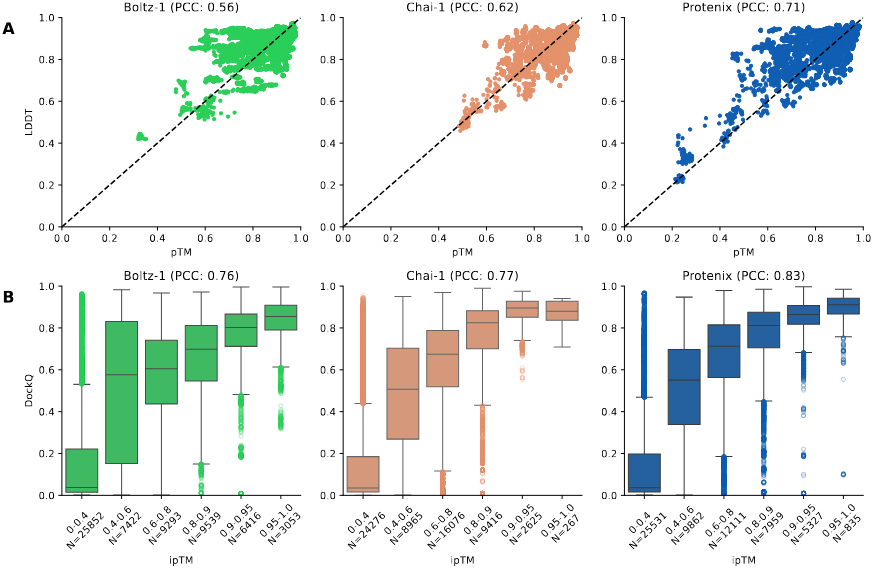
Correlation of self confidence predictions and structural quality. The Pearson correlation coefficients calculated across all 25 samples for each structure. **A**. Monomers. **B**. Interfaces.

**For monomers**, we calculated the PCC between predicted TM-score and LDDT. As shown in Figure 3A, Protenix achieved the highest correlation (PCC = 0.71), followed by Chai-1 (0.62) and Boltz-1 (0.56). This suggests that Protenix’s confidence score is more reliably aligned with structural accuracy in the single-chain setting. **For protein interfaces**, the PCC was computed between interface-predicted TM-score (ipTM) and DockQ. As shown in Figure 3B, Protenix again achieved the strongest correlation (PCC = 0.83), with Boltz-1 and Chai-1 showing comparable but slightly lower values (0.76 and 0.77, respectively).

These results indicate that while all models benefit from confidence-based ranking, Protenix exhibits better consistency between predicted confidence and actual structure quality, especially for multimeric complexes. However, the confidence scores do not fully close the performance gap between oracle and selected predictions (Figure S2). In several cases, high-quality structures were not assigned top confidence, highlighting limitations in current ranking strategies. Future work may benefit from incorporating physics-aware scoring functions or learned reranking mechanisms to further improve selection accuracy.

#### 2.3.3 AF3 vs. Our Protein–Antibody Dataset: A Comparative Study

Within the low-homology RecentPDB subset we assembled, the number of protein–antibody complexes differs markedly from those reported in the AF3 paper. Preliminary inspection suggested that the AF3 list underwent additional filters not specified in their paper. To clarify this discrepancy, we compared the two datasets in detail (Figure 4B).

**Figure 4.**
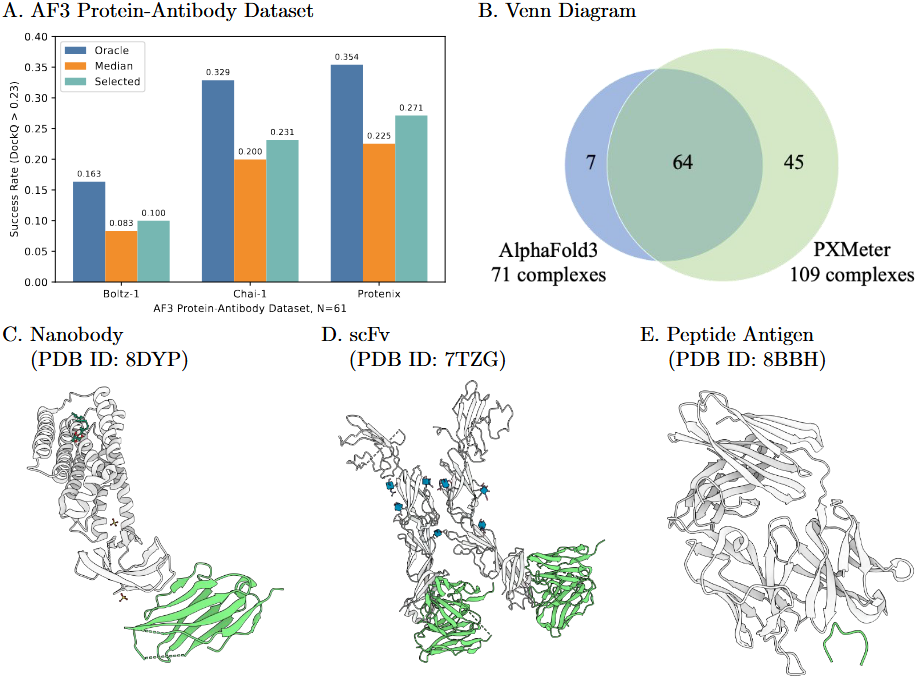
Performance on the AF3 protein-antibody dataset and external complexes.

Of the 64 complexes that the two sets share, all correspond to antigen-binding fragments (Fab). Our dataset contains 45 additional complexes absent from the AF3 list, of which 42 are non-Fab complexes, such as nanobodies (Figure 4C) [19], single-chain variable fragments (scFv, Figure 4D) [20], and Fc fragments (e.g., 8A47). The remaining three complexes are Fab regions. Complex 8BBH was likely excluded because its antigen is a peptide (Figure 4E) [21], whereas the reasons for omitting 7T92 and 7SJN remain unclear.

Conversely, the AF3 protein-antibody dataset contains 7 complexes that we discarded. Three of these were removed for homology to the training set (7UVF, 7SO7, 7R58), three were excluded because less than 30% of the residues were resolved (7ST8, 7RU8, 7UIH), and one complex was excluded because it exceeded the 2560-token limit (7T6X).

To assess the impact of dataset differences on model evaluation, we re-ran benchmarking on the AF3 subset using Boltz-1, Chai-1, and Protenix (4 structures couldn’t be processed at the inference stage by Chai-1 and Boltz-1, see Appendix B). As summarized in Figure 4A, Boltz-1 in particular experienced a pronounced drop in Dock-Q success rate relative to its performance on our curated set, and all three models performed considerably worse than AF3 whose success rate was approximately 0.45 (AF3 paper, Figure 5a).

**Figure 5.**
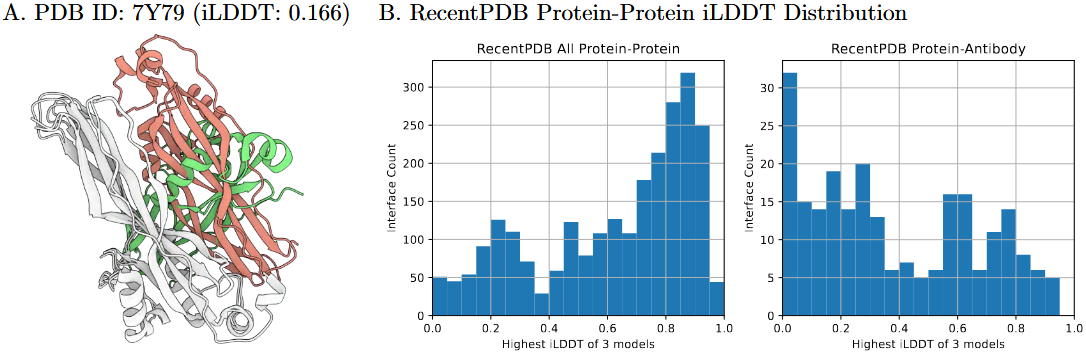
RecentPDB iLDDT Distribution and a Case Study. **A**. The crystal structure of Cry78Aa (shown in gray and green) and the predicted structure with the highest iLDDT score (shown in gray and pink). Chain LDDT: 0.899, 0.900. **B**. Based on the structures predicted by Protenix, Chai-1, and Boltz-1 (5 seeds * 5 samples), the distribution of the highest iLDDT scores for protein-protein interfaces in RecentPDB is presented.

These findings underscore the importance of dataset composition in benchmarking: seemingly subtle differences in antibody formats, antigen types, or resolution thresholds may significantly affect downstream evaluation results and model rankings.

### 2.4 Case Study and Refinement of the RecentPDB Protein-Protein Dataset

#### 2.4.1 Manual Review Reveals Data Quality Issues

To investigate the underlying factors behind the poor performance of problematic cases, we manually reviewed the benchmark subset with low iLDDT scores across all three models (Figure 5B, distribution of iLDDT scores), following the workflow in Figure 1C. Our analysis revealed that weak predictive accuracy is not always due to deficiencies in the models themselves; in many instances, the evaluation data were of questionable quality. For example, the Cry78Aa protein forms a monomer in solution according to the available evidence [22], but its crystal structure (PDB ID: 7Y79) adopts a crystal packing-induced dimer whose contacts do not represent a true biological interface, thus failing to reflect the performance of the models in real biological scenarios. As a control, we randomly sampled a subset of interfaces with iLDDT > 70 and found no systematic data quality issues, suggesting this portion of the dataset is largely reliable. Due to resource constraints, the review was limited to the lowest-scoring interfaces, where errors are most likely to affect performance assessments.

#### 2.4.2 Systematic Identification and Removal of Artifactual Interfaces

We propose that the primary criterion for determining whether an interface should be retained or removed is whether the protein-protein interface is biologically relevant, ideally supported by experimental evidence. The following categories outline the specific circumstances under which interfaces were removed. A detailed list is provided in Appendix F.

- **Crystal Packing:** 57 interfaces were removed from 38 complexes. The removal was based on the criterion of biological relevance in solution. Figure 6A presents an illustrative example. Biochemical and biophysical analysis demonstrated that AcrIIC4 forms a monomer in solution [23], indicating that its dimer observed in the crystal structure was the result of crystal packing. Similar crystal-packing artifacts were observed in the other excluded interfaces.
- **Mislabelled Fusion Protein:** Three interfaces were removed from three complexes where fusion proteins were reported as separate chains. As shown in Figure 6B, structural determination of the human Histamine H3 receptor and E. coli soluble cytochrome b562 involved a fusion of the two proteins [24]. The density map revealed a covalent linkage between the two proteins, rendering this an artifact.
- **Repeat Segments:** 32 interfaces were removed from three complexes. We observed poor performance in predicting fibrous structures formed by multiple repeat segments. Additionally, Bioassembly 1 was modeled with arbitrary repeat-layer assignments [25], as shown in Figure 6C, undermining the validity of the evaluation results.
- **Covalent Link:** Two interfaces were removed from one complex. In complex 7Q5N, an ancient ubiquitin-like protein, ubiquitin-related modifier 1, serves as a sulfur carrier for tRNA thiolation and conjugates with a thioredoxin domain-containing protein [26].
- **Incomplete Modeling:** Two interfaces were removed from one complex. In complex 7FBI, the electron microscopy structure only modeled one-third of the complex. The missing regions prevented accurate reconstruction of the full interface geometry.
- **Duplicate Entry:** Two interfaces were removed from one complex. The authors of complex 7V9X uploaded a more complete version of the complex, identified as 7XJG [27].
- **Other Non-Biological Interaction:** Five interfaces were removed from three complexes. These interactions are non-biological in nature. For instance, in complex 7QNP, lysozyme, which was used for E. coli host lysis, was accidentally co-purified and co-crystallized with protein NA4M4CAII [28].

**Figure 6.**
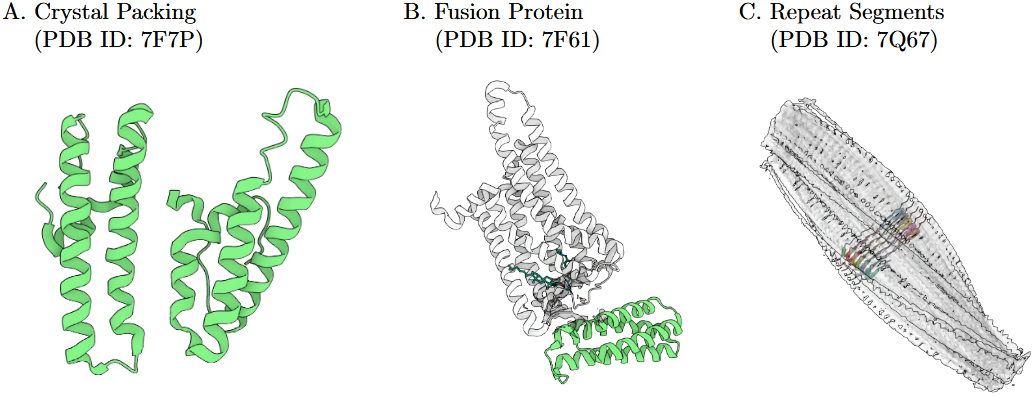
Removed complexes.

#### 2.4.3 Study of Remaining Low-iLDDT Hard Cases

The remaining hard cases comprise 137 experimentally validated protein-protein interfaces (iLDDT < 0.3) originating from 81 distinct complexes, including 82 protein-antibody interfaces from 45 complexes and 55 non-antibody interfaces from 38 complexes.

We found that in many hard cases, antibodies or nanobodies act as structural helper proteins to facilitate structural determination. For instance, in PDB entry 7VAF (Cryo-EM structure of rat NTCP complexed with an antibody Fab fragment [29]), the antibody serves as a fiducial marker to aid in image alignment, with iLDDT values of 0.191 and 0.479 for the two antibody chains. Another example is PDB entry 7YZI (Cryo-EM structure of the adenylate cyclase dimer bound to three copies of a stabilizing nanobody at a resolution of 3.83 Å [30]), where iLDDT values for all protein-antibody interfaces are less than 0.3. The comparison between the predicted structure and ground truth of these two cases is presented in Appendix G.

Some of the hard cases share a common characteristic: they are heterologous assemblies whose subunits originate from different species-often involving host-pathogen interactions, although the exact cause of the model’s underperformance on them remains unclear. Representative examples are provided in Appendix G: the complex between the phage-encoded enolase inhibitor protein from phage *SPO1* and enolase from *Bacillus subtilis* (PDB ID: 7XML) [31] and the interaction between human GSDMB and *Shigella* effector protein IpaH7.8 (PDB ID: 7WJQ) [32]. We speculate that the poor performance observed in these cases may be related to the fact that MSA pairing in AF3-like models is only performed when the different chains originate from the same species.

We also observed that prediction errors manifested at several hierarchical levels of protein architecture. Certain predictions yielded entirely incorrect assemblies—for example, PDB entry 7PUZ. In other instances, while the monomeric structures were similar, the multimeric assemblies differed, as exemplified by 7PV1. Additionally, there were cases where both the monomers and assemblies were reproduced correctly, yet local secondary structures varied—for example, in 7ZGW, the alpha-helix at the binding interface exhibited a translational shift. These three cases are presented in Appendix G.

#### 2.4.4 Performance Update on the Refined Dataset

Through manual review of the low iLDDT protein-protein interfaces, we curated a refined subset of the RecentPDB protein-protein dataset. All models exhibited higher DockQ success rate on this refined subset, as shown in Figure 7, with the most significant gains observed for non-antibody interfaces, as most antibody interfaces were retained unchanged. The rise in metric values reflects the exclusion of artifactual or biologically irrelevant interfaces rather than any change to the models themselves, providing a more accurate picture of their performance on biologically meaningful interactions.

**Figure 7.**
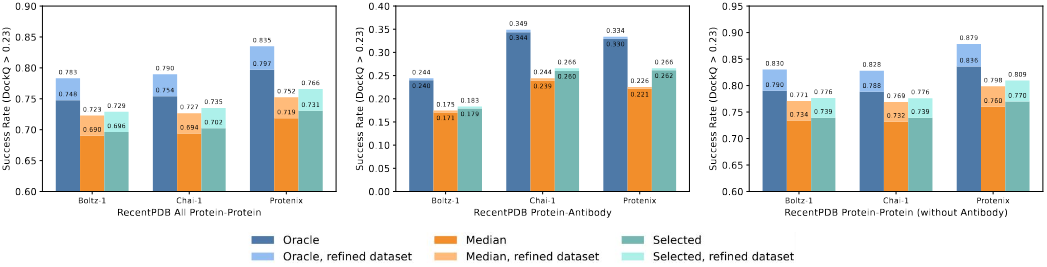
Performance on the refined RecentPDB low-homology protein-protein dataset.

## 3 Discussion

Our benchmarking with PXMeter reveals critical insights into both the capabilities and limitations of current structure prediction methods. By curating challenging evaluation datasets and conducting detailed case studies, we identify recurring failure modes that reflect broader modeling and data issues. These insights could guide the future design of modeling and dataset, helping to improve the robustness and generalization of biomolecular structure prediction.

We observe that current structure prediction models struggle with certain interaction types, especially antibodies or atypical binding modes. In addition, our results show that the models accurately predict individual monomers, but have difficulty assembling them into complex structures, as exemplified by the cases presented in Appendix G, which illustrate incorrect inter-chain orientations or misidentification of the interface. This underscores the need for continued advancements in modeling inter-chain interactions.

As part of the benchmark dataset construction, we manually inspected hundreds of structures and identified multiple artifacts that motivated the exclusion of structures from the evaluation set. The reasons for exclusion included crystal packing, different oligomeric state in crystallization as opposed to solution, fusion of proteins with covalent linkage, arbitrary layer assignments of repeat segments, etc. (Section 2.4). To the best of our knowledge, our benchmark is the first to thoroughly identify these data errors at scale. We provide these findings in Appendix F as a valuable resource for the community. We also emphasize the importance of in-depth case studies of model failures for building a reliable evaluation set and better understanding the applicability of the modeling paradigm. Beyond evaluation set curation, it might be beneficial to extend the data quality filters to training data. Designing automatic and scalable methods for this task remains an ongoing and critical challenge.

Moving forward, addressing these challenges will require improvements in both data pipelines to handle artifacts and expand datasets to cover more diverse biological and synthetic structures, as well as modeling innovations beyond the current AF3 paradigm to better capture complex interactions.

## 4 Conclusion

This work presents a comprehensive framework for evaluating biomolecular structure prediction methods. We introduced PXMeter, an open-source evaluation toolkit designed to enable rigorous and reproducible benchmarking across complex, chain, and interface levels. We curated a diverse and high-quality benchmark dataset by systematically identifying and removing structural artifacts such as crystal packing, repeat segments, and other non-biological interactions. Using PXMeter, we evaluated three state-of-the-art open-source models on our curated dataset, which spans a broad range of biomolecular interaction types. All annotations and filtering criteria are made publicly available to support transparent assessment and reuse.

By providing a unified evaluation toolkit and a standardized benchmark set, this work enables fair comparison across structure prediction methods and facilitates the development of models with high accuracy and generalization.

Our current evaluation dataset and metrics are largely aligned with those used in the AlphaFold 3 study. However, real-world applications in biology and drug discovery involve a wider range of molecular systems and tasks, which require more comprehensive and domain-relevant evaluation. We plan to extend our benchmarks to better capture the diversity of practical scenarios and to support ongoing efforts to improve model robustness and applicability in biomolecular structure modeling.

## 5 Method

### 5.1 How PXMeter Works

PXMeter takes as input the CIF files produced by the structural prediction model ([11–13] and others) and those obtained from the RCSB PDB. It compares the structures within these CIF files and generates a JSON file containing all evaluation metrics.

Given the two CIF files, PXMeter first calculates the sequence identity for each entity. Then it matches the entities in descending order of sequence identity to establish entity pair correspondence. This mapping step is robust to sequence discrepancies, such as missing residues in the model, thereby enabling fair evaluation even when the two sequences are not identical. Within each matched entity pair, the corresponding chains from both the reference and the model structures are selected, ensuring that the intersection of residues is identified and that all atoms are aligned sequentially. For the model structure, equivalent chains and atoms within an entity are permuted to better align the prediction with the true structure, as detailed in the Appendix A).

PXMeter currently supports a variety of evaluation metrics, including the LDDT for complex, chain, and interface levels, DockQ for interface evaluation, as well as pocket-aligned RMSD and PoseBusters plausibility checks for specific ligands. The DockQ metric is calculated using the DockQ V2 [33] Python API, with PXMeter’s chain mapping results provided as input for precise assessment. Similarly, PoseBusters plausibility checks are performed through the PoseBusters [8] Python API, utilizing the mapped ligand and protein files supplied by PXMeter for evaluation.

For nucleic acid entities, the LDDT inclusion radius is set to 30 Å due to their larger size, while other entities use a 15 Å radius, following the setting in the AF3 paper.

### 5.2 Dataset Construction

#### RecentPDB

This dataset was curated from CIF files downloaded from the RCSB PDB on 2024-05-22, with extracted information utilized for filtering purposes. Based on the RecentPDB construction method from the AF3 paper [1], a dataset is curated for evaluation purposes. Water molecules and hydrogen atoms were removed from all structures, after which the structures were expanded to Assembly 1 as recorded in the CIF files. After applying all the filters described in this section, 8,865 structures were retained.

Filtering of complex:

- From a total of 220,113 PDB entries, 10,172 entries with release dates between 2022-05-01 and 2023-01-12 were selected.
- Filtering to 9,944 non-NMR entries.
- Filtering to 9,637 entries with resolutions better than 4.5 Å.
- Select 8,875 entries where the number of tokens was less than 5,120 under AF3’s tokenization scheme.

Filtering of chain:

- A polymer chain was retained only when the “_entity_poly.type” field recorded in the CIF file was annotated as “polypeptide(L)”, “polydeoxyribonucleotide”, or “polyribonucleotide”. Other polymer types, such as “polydeoxyribonucleotide/polyribonucleotide hybrid” or “peptide nucleic acid”, are uncommon and unsupported by some of the current structure prediction models, and were therefore excluded. Non-polymer chains were all categorized as “ligand” and retained.
- Chains with all residues unknown were removed.
- If the number of resolved residues in a chain is less than four or less than 30% of the total sequence length, the chain was removed. This step reduced the number of RecentPDB entries by ten, down to 8,865.

#### RecentPDB Protein Low Homology Subset

To evaluate generalization to unseen sequences, we constructed a low-homology subset based on sequence identity to the model training set, using a cutoff date of 2021-09-30. Each structure in this subset was used in full as input for structure prediction. Evaluation metrics, however, were computed only on protein chains and protein-protein interfaces with low sequence identity to training data. Within the biological Assembly 1, an interface (a chain pair) was defined whenever any pair of heavy atoms from two distinct chains were *≤*5 Å apart. After applying the following filters, 1569 entries remained.

Filtering of complex:

- Selected entries where the number of tokens was less than 2,560.

Filtering of chain and interface:

- Peptide chain: Filtered out peptide chains, where a peptide was defined as a protein chain with fewer than 16 residues.
- Protein chain: Kept only chains with sequence identity to the training set below 40%.
- Peptide-protein interface: The protein chain must have less than 40% sequence identity to any sequence in the training set.
- Peptide-peptide interface: Filtered out.
- Protein-protein interface: If both chains had sequence identity greater than 40% and in the same complex in the training set, then the interface was filtered out.

We use hmmsearch [34] to identify training set sequences that are similar to those in the test set. These sequences are then aligned using Kalign [35], and sequence identity is computed. Our approach follows the template search procedure described in Section 2.4 of the AF3 paper’s Supplementary Information.

Ultimately, 1,569 entries were selected, comprising 2,475 intra-protein chains and 2,773 protein-protein interfaces. During the evaluation process using this dataset, we manually reviewed certain chains and interfaces where the models performed poorly. As a result, we excluded some structures that were deemed unsuitable for evaluation. For more details, please refer to Figure 1B and Section 2.4.

Clustering:

Individual polymer chains were clustered at a 40% sequence identity for proteins with more than 9 residues and 100% sequence identity for protein with less than or equal to 9 residues.

- Protein chain (> 9 residues): The cluster ID is obtained directly from the sequence clustering file provided by RCSB PDB (https://cdn.rcsb.org/resources/sequence/clusters/clusters-by-entity-40. txt, downloaded at 2024-06-30).
- Protein chain (*≤* 9 residues): The cluster ID is simply the string of its sequence
- Protein-Protein interface: The cluster ID is (protein1 cluster, protein2 cluster)

Protein monomer and protein-antibody subset:

In the evaluation, protein monomers and protein-antibody interfaces are identified from the RecentPDB Protein Low Homology Subset, allowing separate assessment of the model performance on these data categories.

- Protein monomer: A chain is considered a protein monomer if Assembly 1 of the entry contains only one protein chain and no other polymer chains. There are a total of 508 protein monomer entries here.
- Protein-antibody interface: An interface is classified as a protein-antibody interface when one protein chain belongs to any of the two largest clusters of the RCSB PDB sequence cluster file, whereas the other protein chain is not part of these clusters. A total of 109 entries are selected, encompassing 246 protein-antibody interfaces.

#### AF3-AB Subset

AF3 [1] released an protein-antibody subset (AF3-AB) that differs from the one we have curated. We also incorporated in our evaluation this AF3-AB subset retrieved directly from the official GitHub repository [36].

#### dsDNA-Protein, RNA-Protein Subset

A nucleic acid evaluation subset was derived from the RecentPDB dataset. We began with the PDB IDs listed in Supplementary Table 14 of the AF3 paper, and then applied a series of filters as follows:

- The maximum number of residues in a structure was limited to 1000 to maintain consistency with the criteria used in the AF3 paper.
- The dataset was restricted to include only those PDB entries with one RNA chain plus one protein chain, or two DNA chains (double stranded DNA) plus one protein chain. Complexes containing both DNA and RNA were excluded.
- DNA sequences bearing modified nucleotides were removed.

Ultimately, the filtering yielded 38 dsDNA-Protein interfaces and 25 RNA-Protein interfaces for evaluation. Although these totals matched the counts reported in the AF3 paper, we cannot confirm that the PDB IDs are identical to those used in their study.

#### PoseBusters

To assess protein–ligand modeling performance, we adopted the PoseBusters V2 [8] benchmark set. For the exceptionally large complex in PDB entry 8F4J, we retained only protein chains located within 20 Å of the ligand of interest, following the same approach described in the AF3 paper. All evaluations were carried out on the asymmetric unit. For each complex, the ligand metadata were parsed from the RCSB PDB CIF files. When multiple symmetry related copies of the ligand were present, only the designated instance with the correct alternative location was utilized for RMSD computation, ensuring that every complex was represented by a single, unambiguous ligand model.

### 5.3 Reproducible Evaluation

#### 5.3.1 Setting of Model Inference

Protenix (v 0.5.0) [13], Chai-1 (v 0.6.1) [11], and Boltz-1 (v 0.4.1) [12] were used for evaluation. Each model was run with five random seeds, generating five samples per seed. Inference was performed using 10 recycling iterations and 200 diffusion steps.

All input structures were preprocessed to remove crystallization aids. Covalent bonds between the ligand and the polymer were retained. Glycan chains were handled in a model-specific mannar: for Chai-1, carbonhydrate chains were supplied using the built-in glycan syntax; for Protenix, glycans were represented as a sequence of multiple CCD codes linked by covalent bonds; the current release of Boltz-1 lacks native support for glycan objects, so all glycan chains were removed from its input files.

The same MSA file was supplied to all models to provide uniform sequence information. MSA input was restricted to protein chains only. We searched MSAs using MMSEQS2 [37] and ColabFold [38] MSA pipeline, and used MSAs from the Uniref100 [39] database for pairing (species are identified with taxonomy IDs).

#### 5.3.2 Aggregation of Metrics

Metrics were computed with PXMeter for each of the 25 samples generated by the three models and a single representative value was chosen for every complex, chain and interface. Because the optimal sample can vary across structural elements, the representative metrics for distinct interfaces or chains within the same complex may originate from different samples. The resulting values were subsequently averaged within each evaluation subset. For the chains or interfaces associated with a cluster ID, the metrics were first averaged across all members of the cluster to yield a cluster level mean, and these cluster means were then used in the subset level aggregation.

## 6 Code and Data Availability

Source code and the curated dataset are available at https://github.com/bytedance/PXMeter.

## Appendix

### A Chain and Atom Permutation

Our chain permutation procedure builds upon the heuristic algorithms implemented in AF3 [1] and AF2.3 [40]. For each predicted complex, the chain indices are rearranged to minimize the overall RMSD with the reference structure. Initially, anchor chains are designated in both the predicted and ground truth structures - preferably polymer chains containing at least four resolved residues. Subsequently, all potential anchor pair combinations are enumerated, with the remaining chains being matched greedily for each pair. The permutation that results in the lowest global RMSD is retained.

The atom permutation procedure begins by constructing, for each residue (or ligand; hereafter residue denotes either), a graph whose vertices represent the atoms and edges encode covalent connectivity, irrespective of bond order. Graph-isomorphism analysis is then applied to enumerate all exchangeable atom mappings, with at most 1,000 permutations retained per residue. Each residue is subsequently matched individually, selecting the atom mapping that minimizes its intra-residue RMSD.

### B Overview of Successfully Inferred Entries Count

|  | RecentPDB Protein LowH | PoseBusters | AF3-AB | dsDNA-Protein | RNA-Protein |
| --- | --- | --- | --- | --- | --- |
| **Total** | 1569 | 308 | 71 | 38 | 25 |
| Protenix | 1566 | 308 | 71 | 38 | 25 |
| Chai-1 | 1478 | 299 | 67 | 38 | 25 |
| Boltz-1 | 1527 | 301 | 67 | 38 | 25 |

There were three entries—7Z5O, 7T2Q, 7×1R—that failed Protenix inference due to the presence of “U” in their protein sequences.

The inference failures of Chai-1 and Boltz-1 stemmed from two primary causes: Chai-1’s 2048-token limit and Boltz-1’s running out of GPU memory.

All structures inferred by the model were successfully evaluated with PXMeter without any failures.

### Comparison of Metrics Calculated by PXMeter and OpenStructure

**Figure S1.**
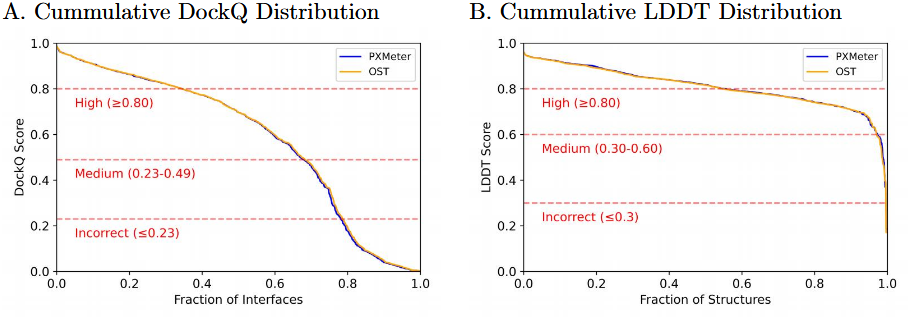
Cummulative DockQ/LDDT Distribution of PXMeter and OST. **A**. Cumulative distribution of DockQ scores on protein-protein interfaces. **B**. Cumulative distribution for complex LDDT scores.

OpenStructure (OST) [4] is a widely used evaluation tool in the field of structural prediction. To validate PXMeter, we compared its DockQ and complex LDDT scores (OST does not provide interface LDDT scores) with those generated by OST. The two sets of metrics agreed closely, confirming the numerical fidelity of PXMeter’s implementations.

The 585 test entries were sourced from the RecentPDB protein low homology subset, focusing on instances where the asymmetric unit is identical to Assembly 1. During the LDDT test, we removed the non-polymer parts from the complexes and set the PXMeter nucleic acid’s LDDT inclusion radius to 15 Å to align as closely as possible with OST’s settings. The version of OST used for testing was 2.8.0, with the testing command being

~~~
ost compare - structures -r [ ref_cif] -m [ model_cif] -- dockq - rna -- lddt -- lddt - no - stereochecks.
~~~

### D Examples of Protenix Predicted Models Compared with Crystal Structures

**Figure S2.**
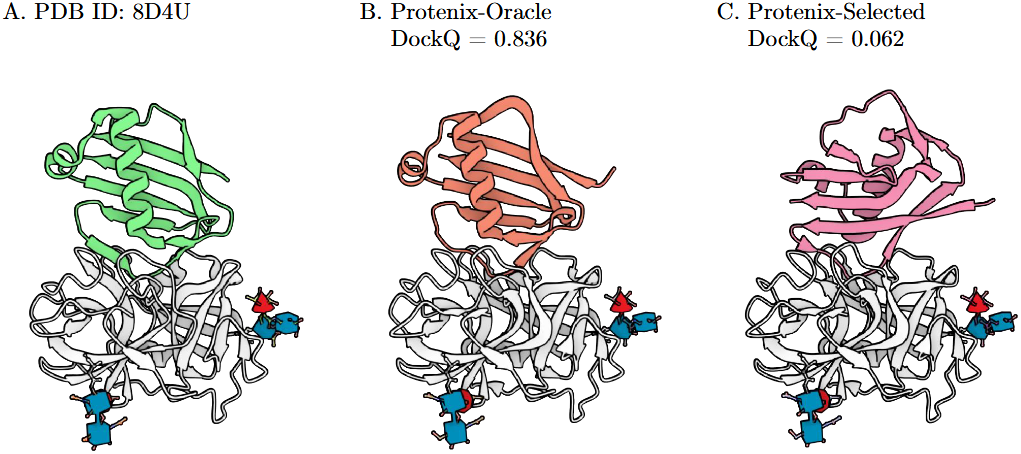
The Protenix prediction and crystal structure of neutrophil elastase inhibited by Eap2 from S. aureus. The Protenix model was able to sample near-native structure as candidate but the confidence score did not reliably identify the correct binding pose.

**Figure S3.**
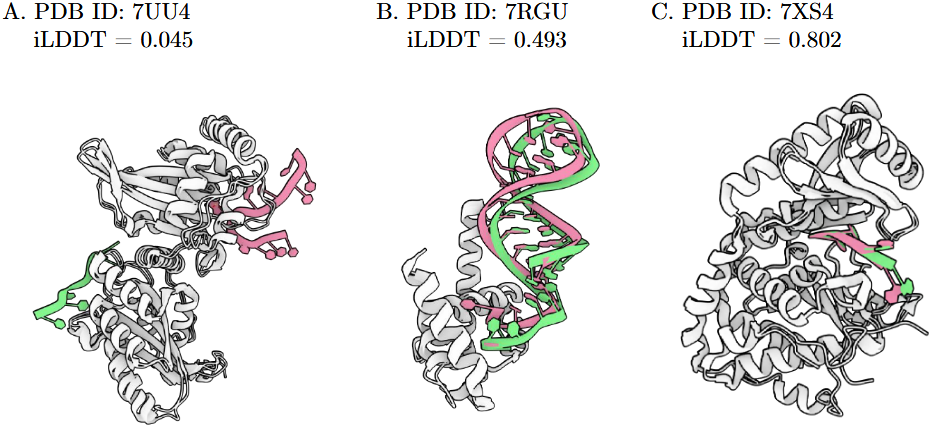
The Protenix-selected prediction (pink) and crystal structures (green) of sampled protein-rna complexes. Low iLDDT value indicates a completely wrong interaction motif. Moderate iLDDT value indicates a correct binding site but slightly deviated interaction motif. High iLDDT value indicates a near native structure.

### Additional Model Performance Comparison

**Figure S4.**
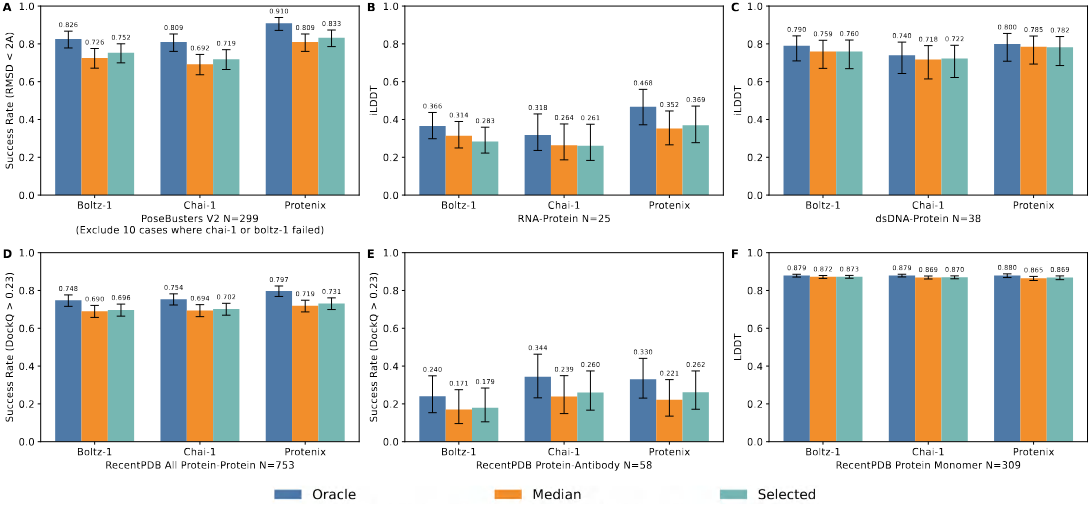
Performance comparison of the Boltz-1, Chai-1, and Protenix models on the curated benchmark datasets. The plot uses the same data as Figure 2, but features a consistent Y-axis for more precise comparison and incorporates error bars. The error bars represent 95% confidence intervals, derived from exact binomial distribution for PoseBusters V2 and from 10,000 bootstrap resamples for all others.

**Figure S5.**
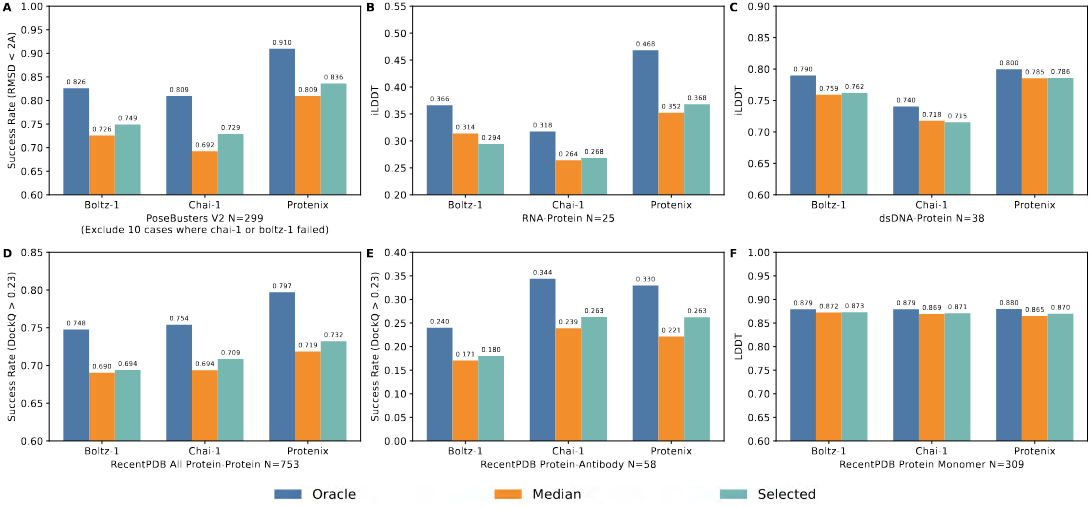
Performance comparison of the Boltz-1, Chai-1, and Protenix models on the curated benchmark datasets. The <u>Selected</u> score in this figure reports metrics for the top-ranked chain/interface structure according to the model’s **chain pTM** for protein monomer and **chain-pair ipTM** score for others.

**Figure S6.**
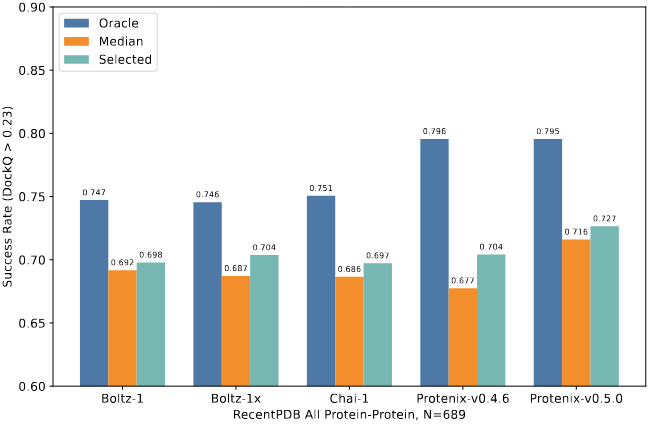
Performance of the Boltz-1x and Protenix-v0.4.6 models on the RecentPDB Low Homology Protein-Protein Interfaces. During inference, Boltz-1x (v1.0.0) experienced more “out of GPU memory” errors than Boltz-1 (v0.4.1). However, for successful inference entries, both models exhibited similar performance. Compared to the previous version v0.4.6, Protenix-v0.5.0 demonstrates improvements in the median and selected score.

### F Table: Protein-Protein Interfaces Removed After Manual Curation

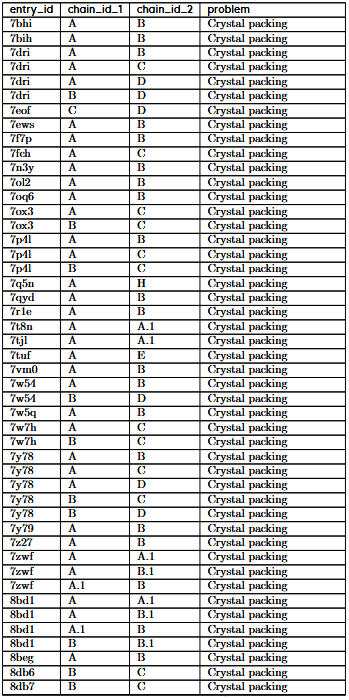
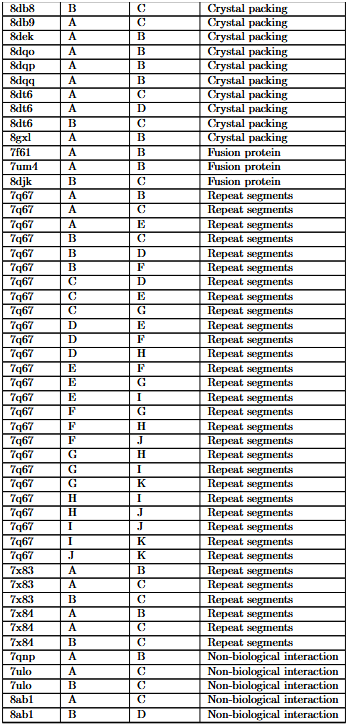
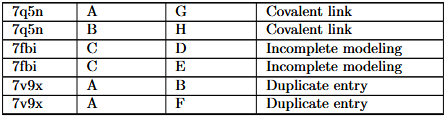

### G Examples of Hard Cases

The table presents hard cases from the RecentPDB dataset (experimentally validated protein-protein interfaces with iLDDT < 0.3). The visualized structure corresponds to the highest-complex-LDDT prediction among the three model outputs.

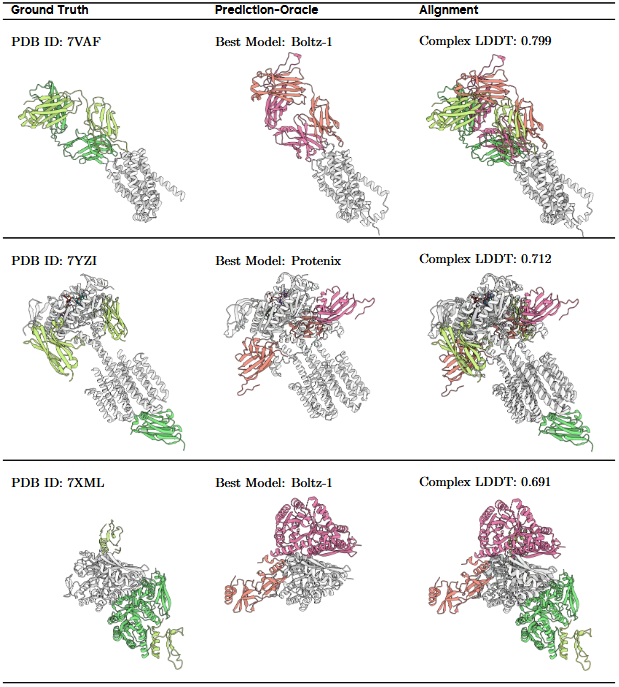
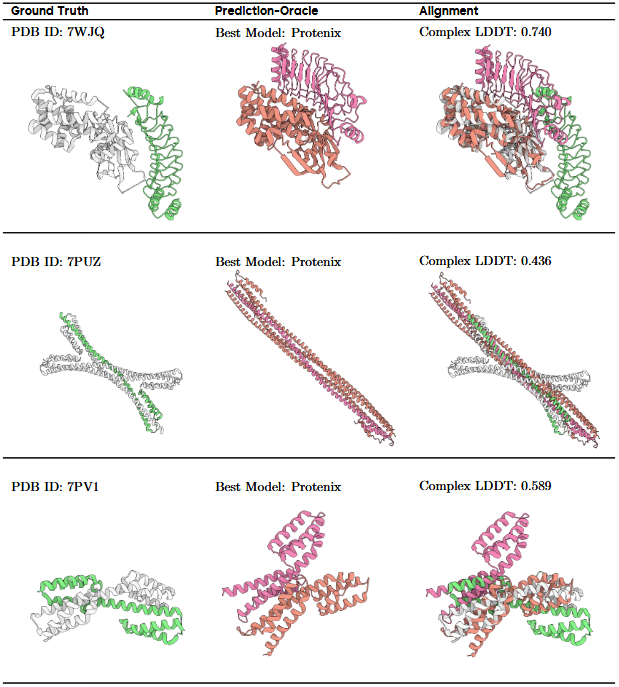
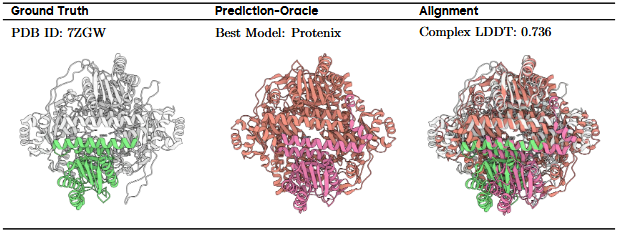

